# Integrated single-nucleotide and structural variation signatures of DNA-repair deficient human cancers

**DOI:** 10.1101/267500

**Authors:** Tyler Funnell, Allen Zhang, Yu-Jia Shiah, Diljot Grewal, Robert Lesurf, Steven McKinney, Ali Bashashati, Yi Kan Wang, Paul C. Boutros, Sohrab P. Shah

**Author notes:** Correspondence and requests for materials should be addressed to S.P.S.

## Abstract

Mutation signatures in cancer genomes reflect endogenous and exogenous mutational processes, offering insights into tumour etiology, features for prognostic and biologic stratification and vulnerabilities to be exploited therapeutically. We present a novel machine learning formalism for improved signature inference, based on multi-modal correlated topic models (MMCTM) which can at once infer signatures from both single nucleotide and structural variation counts derived from cancer genome sequencing data. We exemplify the utility of our approach on two hormone driven, DNA repair deficient cancers: breast and ovary (n=755 cases total). Our results illuminate a new age-associated structural variation signature in breast cancer, and an independently identified substructure within homologous recombination deficient (HRD) tumours in breast and ovarian cancer. Together, our study emphasizes the importance of integrating multiple mutation modes for signature discovery and patient stratification, with biological and clinical implications for DNA repair deficient cancers.

---

Patterns of mutation in cancer genomes reflect both endogenous and exogenous mutagenic processes^1^, allowing inference of causative mechanisms, prognostic associations^2^, and clinically actionable^3–6^vulnerabilities in tumors. Many mutational processes leave distinct genomic “footprints”, measurable via nucleotide substitution patterns^1^, localised mutation densities, and patterns of structural variation. As such, each mutagenic source (whether exogenous or endogenous) changes DNA in a characteristic manner, at genomic locations with preferred chemical and structural characteristics. Exogenous insults such as ultra-violet radiation and tobacco smoke associated mutagens (*e.g.* benzo[a]pyrene) induce single nucleotide variants (SNVs) with characteristic C→T (at CC or TC dinucleotides)^7^ and C→A mutation patterns^8^, respectively; endogenous APOBEC3A mediates enzymatic 5-methylcytosine deamination, resulting in C→T substitution patterns at TC dinucleotides^7^.

Cancer cells can also acquire an endogenous mutator phenotype, accumulating large numbers of mutations^7^ due to DNA repair deficiencies. Defective DNA repair processes induce both point mutations and structural variations^9^, and include several mechanistic classes such as mismatch repair deficiency, homologous recombination deficiency, microhomology mediated end-joining, and breakage fusion bridge processes. Defective DNA repair has been exploited in therapeutic regimes, including immune checkpoint blockade for mismatch repair deficiency^6^, and synthetic lethal approaches for homologous recombination deficiency^4,5^, underscoring their clinical importance.

Both point mutation signatures^10^ and structural variation signatures^11^ have been studied extensively as independent features of cancer genomes, mostly through non-negative matrix factorization (NMF) approaches^1,3,12–15^. As increasing numbers of whole genomes are generated from tumors in international consortia and focused investigator research, the need for robust signature inference methods is acute. Additional computational methods have been proposed^16–19^, however no approaches jointly infer signatures from *both* point mutation and structural variations. We contend that systematic, integrative analysis of point mutation and structural variation processes enhances ability to exploit signatures for subgroup discovery, prognostic and therapeutic stratification, clinical prediction, and driver gene association.

Latent dirichlet allocation (LDA)^20^, a popular and effective approach for natural language document analysis, is well suited to the task of mutation signature inference. Appropriate conceptual mappings applied to mutation signature analysis can be described as follows: signatures (topics) are represented as distributions over a mutation (word) vocabulary, and sample mutation catalogues are represented as distributions over mutation signatures. In this paper we introduce the correlated topic model (CTM)^21^, an extension of LDA which incorporates signature correlation, and a multi-modal correlated topic model (MMCTM)^22^ which jointly infers signatures from multiple mutation types, such as SNVs and SVs. Signature activities can be correlated among some groups of patients, motivating the use of this class of methods. For example, homologous recombination deficiency induces patterns of both SNVs and SVs in breast^13^ and high grade serous ovarian cancers^2^. We show how integrating SNV and SV count distributions improves inference of signatures relative to NMF and standard topic modeling methods.

Motivated by the need to better understand mutation signatures in the context of DNA repair deficiency, we applied the MMCTM to SNV and SV somatic mutations derived from publically available breast^13^ and ovarian^2^ whole genomes (755 cases total), performing joint statistical inference of signatures. Our results reveal correlated topic models as an important analytic advance over standard approaches. Rigorous benchmarking over mutation signatures inferred from previously published mutation corpora was used to establish metrics for comparison. In addition, we report novel strata using MMCTM-derived signatures, including patient groups exhibiting combined whole genome SNV and SV signature profiles from breast and ovary cancers. We also automatically recovered BRCA1-like and BRCA2-like homologous recombination repair deficient breast and ovarian cancers, where the tumors bearing the well known SNV HRD signature were reproducibly split on the basis of SVs. We further uncovered prognostically relevant strata in ovarian cancer, identifying important patient subgroups for further clinical and biologic study. In aggregate, our study reveals the importance of simultaneously considering multiple classes of genomic disruption as a route to expanding mutation signature discovery, and their downstream impact on novel stratification across human cancers.

## Results

### Datasets and feature construction

We studied mutation signatures in 560 breast^13^ and 195 ovarian^2,23^cancer genomes (Supplementary Table 1). Each dataset was analyzed separately to avoid biases from differences in sample sequencing, data-processing or annotation.

We constructed SNV features using the 6 types of pyrimidine-centric substitutions (C→A, C→G, C→T, T→A, T→C, C→G), and their flanking nucleotides. SNV signature analyses have traditionally focused on the variant and two flanking nucleotides (one 5′ and one 3′ to the variant)^10,24^. Here, we used four flanking nucleotides (two 5′ and two 3′ to the variant) for identifying SNV context bias. We defined SV features by rearrangement type (deletion, tandem duplication, inversion, fold-back inversion (FBI), translocation), number of homologous nucleotides around the breakpoints (0-1, 2-5, >5), and breakpoint distance (<10kbp, 10-100kbp, 100kb-1Mbp, 1-10Mbp, >10Mbp, except for translocations).

We implemented several “dependent” feature methods (LDA, CTM, MMCTM, Supplementary Table 2) which, like NMF, accept counts of mutation “words” that incorporate the mutation type itself, and contextual information. For example, a C→T SNV with an upstream A and downstream G can be represented as the item “A[C→T]G”. In addition, we implemented “independent” feature models^16^ (ILDA, ICTM, IMMCTM, Fig. 1a, Supplementary Table 2) which treat each mutation as a collection of features, rather than as a single vocabulary item. That is, one feature for the mutation itself (say, C→T), and features for each piece of contextual information (*e.g.* 5′ A and 3′ G in the previous example). Assuming 6 SNV types, and 4 flanking nucleotides (two 5′ and two 3′ to the variant), the number of features is reduced from 6 * 4^4^ = 1536 for dependent feature models, to 6 + 4 * 4 = 22.

**Figure 1:**
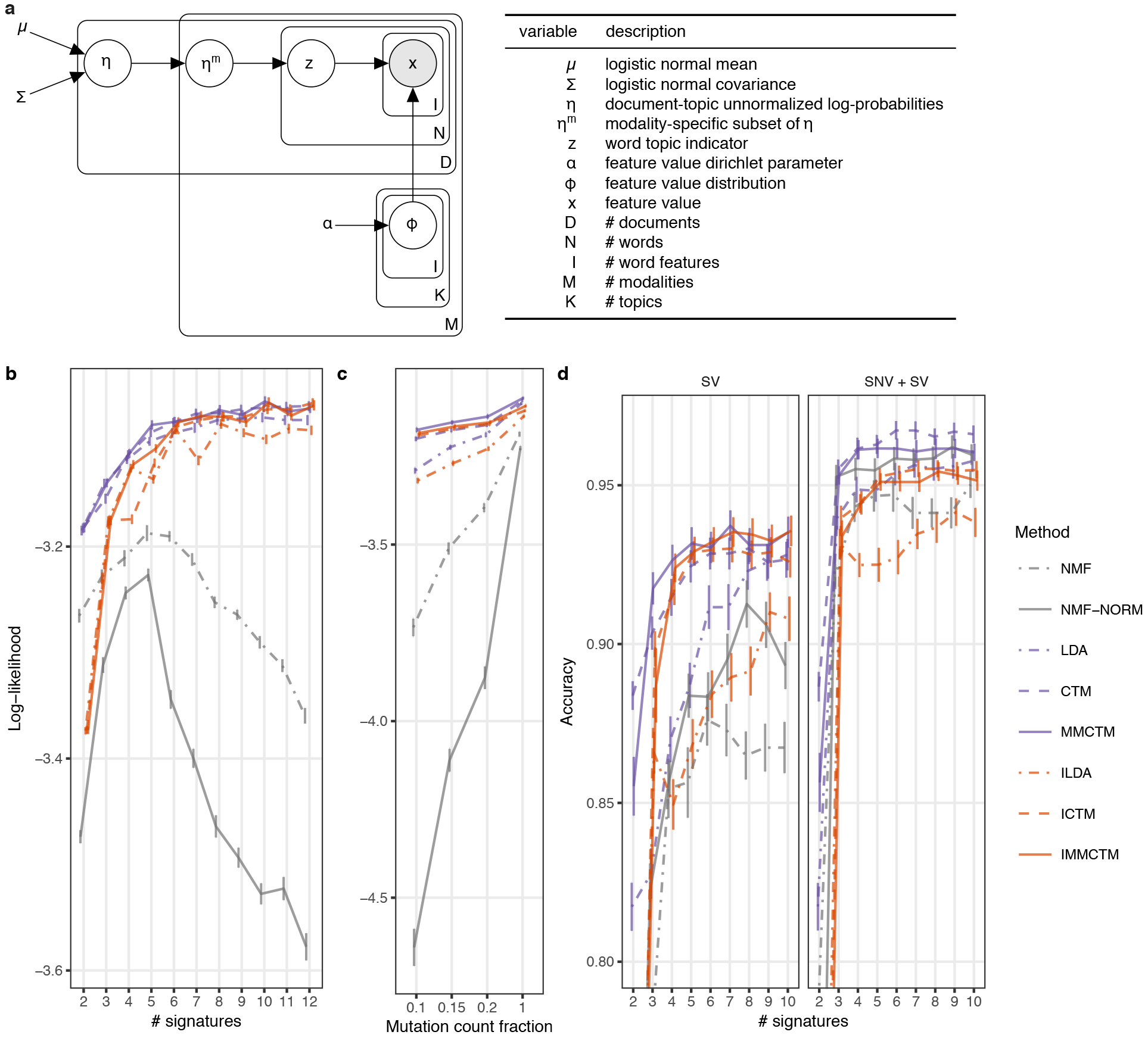
IMMCTM graphical model and comparison of methods using the 560 breast cancer dataset. a IMMCTM graphical model and variable descriptions. SV signature log likelihood means ± standard error for: **b** 2-12 signatures, and **c** a range of SV mutation count fractions. **d** Logistic regression accuracy means ± standard error for predicting HRD labels using per-sample signature mixture proportions across a range of 2-10 signatures (each for SNV and SV). Accuracy is displayed for training with SV (top) and both SNV and SV (bottom) signature proportions. NMF: applied to raw counts, NMF-norm: applied to normalized counts. Vertices and error bars for all plots are dodged slightly to reduce overplotting.

Model specifications and implementation details for all methods are provided in the Supplementary Materials and Methods sections.

### Correlated topic models provide improved signature inference

We compared NMF to LDA, CTM, MMCTM, and the independent feature models ILDA, ICTM, and IMMCTM (Fig. 1a). As NMF is commonly given normalized mutation counts, we also included a normalised input to NMF (NMF-NORM). We performed 5-fold cross validation, repeated 10 times, on the breast cancer dataset described above. For each comparison, we fit SNV and SV signatures to four folds and computed the average per-mutation predictive log-likelihood on the held out fold (112 samples). We split mutation counts from each test sample into two parts, inferred sample-signature activities with one portion, and computed log-likelihood values with the other portion. This evaluation procedure required mutation signatures and sample-signature activities from each method.

We first compared performance as a function of the number of signatures, fitting models over a range of 2-12 SNV and SV signatures (Fig. 1b, Supplementary Fig. 1a, Supplementary Data 1). For SV signatures, NMF performance degraded with >5 signatures, while the probabilistic topic models’ performance improved until a plateau was reached. The dependent and independent feature correlated models performed comparably at inferring SV signatures, while NMF-NORM performed worse than NMF. For SNV signatures, LDA, CTM, and MMCTM all performed best. ILDA, ICTM, and IMMCTM performed much worse than other methods, but their performance continually improved over the range of tested signature counts, eventually matching NMF.

Correlated topic models performed better than their non-correlated analogues at inferring SV signatures, possibly due to relatively low input counts for SNV and SV features. To explore this further, we compared performance over a range of mutation count fractions (Fig. 1c, Supplementary Fig. 1b, Supplementary Data 2). With fewer SV counts, MMCTM outperformed CTM, which outperformed LDA. When subsetting SNV counts, LDA, CTM, and MMCTM performed roughly equally until only 1% of mutation counts were retained, at which point LDA performance became worse than the CTM and MMCTM. Importantly, correlated topic models were the least affected by reducing mutation counts, whereas NMF exhibited the worst performance decline, indicating that correlated models were in general more robust to data sparsity.

We next compared the quality of patient stratification, where the input features were computed by the respective methods (Fig. 1d, Supplementary Fig. 1c, Supplementary Data 3). We trained each method 10 times with random initializations on the full breast cancer dataset. We then trained a logistic regression classifier with the per-sample signature activities from each run, and published HRDetect-derived HRD labels^3^. HRD prediction accuracy scores were computed using 5-fold cross-validation. When the classifier was trained on only SNV signature activities, the CTM and MMCTM performed equally well. NMF-NORM generally did at least as well as the CTM and MMCTM in this comparison, but NMF with raw counts performed worse. With SV signature activities, the correlated topic models converged to a similar performance, and generally provided better average accuracy than the other methods. When the classifier was trained on both SNV and SV signature activities, the CTM and MMCTM performed better than other methods, and the CTM performed somewhat better for signatures ≥ 6. NMF-NORM also had slightly worse, but comparable performance to the MMCTM, in this comparison.

Overall, correlated topic models produced superior predictive mutation signature distributions and low-dimensional representations of samples. This was especially true when each sample had few mutations, as for SVs. This may suggest that they would perform better with exome data than other types of models. We found similar patterns in log-likelihood comparisons using the smaller ovarian cancer dataset, except we detected no differences between the CTM and MMCTM, and the independent feature models performed best with 1% SNV counts (Supplementary Fig. 2). Performance of probabilistic topic models was stable across a range of topic hyperparameter values (Supplementary Fig. 1d).

### Integrated SNV and SV signatures in breast cancer

We next analysed the 560 breast cancer genomes^13^ with the MMCTM (Supplementary Fig. 3a) for stratification analysis. We simultaneously fit 8 SNV and 8 SV signatures to counts of SNVs and SVs (Fig. 2a,b, Supplementary Fig. 4, Supplementary Data 4, see Methods for signature count selection). We found signatures similar to those identified previously^13^, including the age-related (SNV-7, COSMIC 1, Supplementary Fig. 5), APOBEC (SNV-4 & SNV-6, COSMIC 2 & 13), deletion (SV-5), and tandem duplication (SV-4, SV-6, SV-8) signatures.

**Figure 2:**
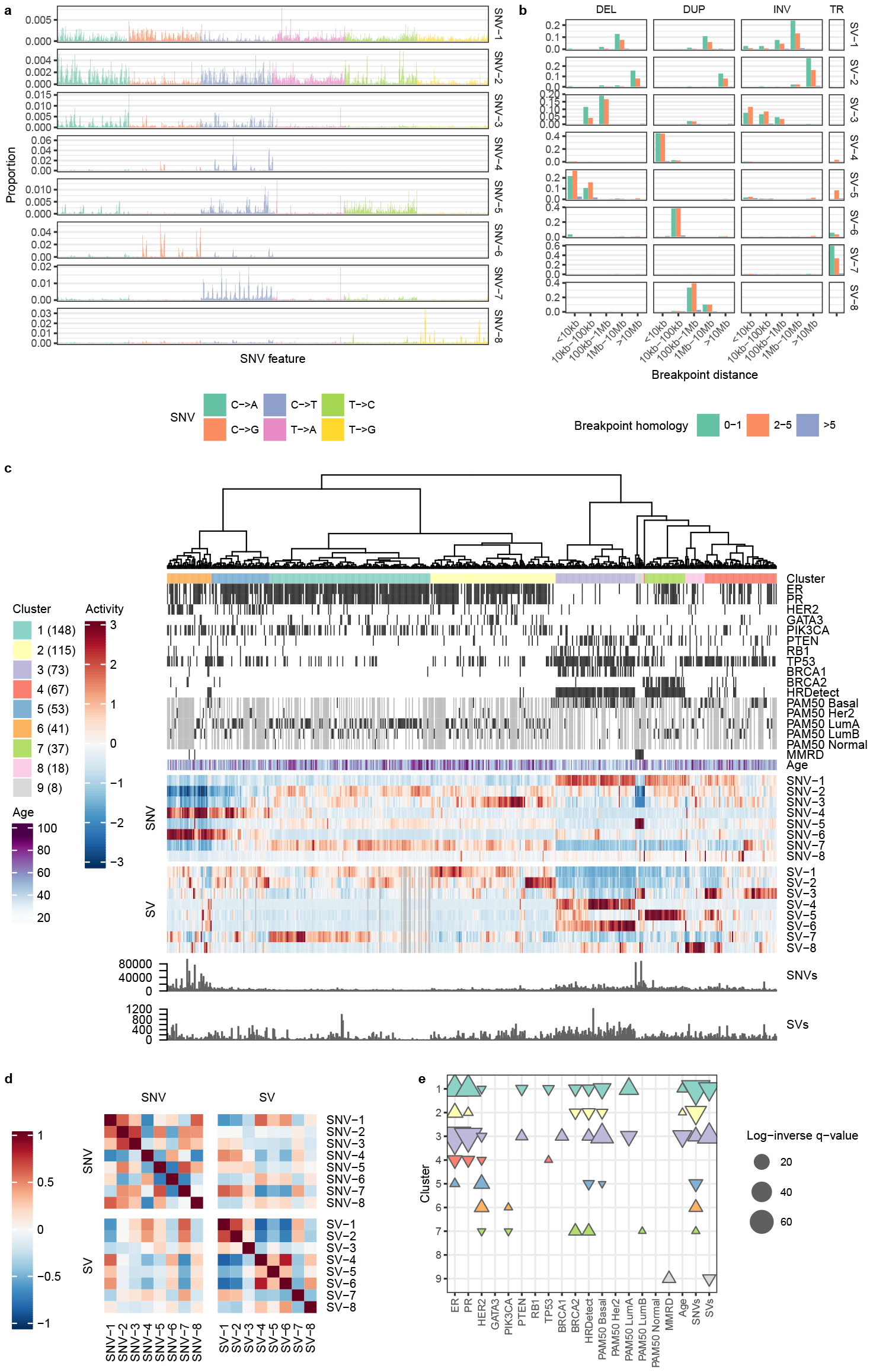
BRCA-EU mutation signature analysis. a SNV mutation signatures. SNVs are organized according to the SNV type (color). Within each type, SNVs are further organized into the pattern of flanking nucleotides (AA-AA, AA-AC, … TT-TG, TT-TT). **b** SV mutation signatures. SVs are grouped by type (DEL: deletion, DUP: tandem duplication, INV: inversion, TR: translocation). **c** Heatmap of relative signature activities in BRCA-EU samples. Each heatmap column represents a single case, and is composed of the proportions of SNV and SV signatures output from the MMCTM model. The values for each signature (row) have been standardized. Heatmap display has been truncated to ±3. Cases have been heirarchically clustered according to these transformed signature prevalences and cluster labels are indicated with colors underneath the dendrogram. The number of cases in each cluster is indicated in paretheses in the cluster legend. ER, PR, and HER2 positive status, gene driver mutation status, HRDetect prediction, and MMRD status is indicated with black bars. Grey cells represent missing data for annotation tracks. Samples with zero mutations for a mutation type also have greyed mutation signature activity cells. **d** Correlation heatmap between SNV and SV signatures. **e** Annotation associations for sample clusters. Upward- and downward-pointing triangles indicate enrichment and depletion, respectively. Colors correspond to cluster colors indicated in the heatmap.

Some signatures were more likely to co-occur in the same tumour, possibly reflecting common etiology. For example, the two APOBEC signatures were positively correlated (Pearson’s r=0.38) (Fig. 2d, Supplementary Data 5), and the HRD SNV signature was positively correlated with the small tandem duplication signature (r=0.59), as expected. The age-related signature (SNV-7) was positively correlated with signatures of intrachromosomal SVs 1-10Mbp (SV-1, r=0.6) and >10Mbp in size (SV-2, r=0.51).

We next performed unsupervised clustering over tumours on joint per-tumour SNV and SV signature activities (Fig. 2c, Supplementary Figs. 3b, 6, Supplementary Data 6, 7, see Methods). The resulting 9 groups included two (clusters 1 & 2, n=148 & 115) enriched for the age-associated signature (SNV-7, see Supplementary Fig. 7a, Supplementary Data 8 for significant cluster-signature associations). Cluster 2 was distinguished from cluster 1 in part due to enrichment of SNV-3. Cluster 1 had the highest relative activity of SV-7 (translocations), while both clusters 1 & 2 had enriched activity of large intra-chromosomal rearrangements (SV-1 & SV-2), especially cluster 2. While SNV-7 was most correlated with age (r=0.23), SV-1 was second most correlated (r=0.17). Cluster 1 was associated with Luminal A cancers, and both clusters 1 and 2 contain tumours from generally older patients with relatively fewer SNVs (see Fig. 2e, Supplementary Data 9 for significant cluster-annotation associations). This implies that older patients may be more likely to have accumulated SVs in their cancers’ etiology as function of background rates, indicating a putative SV-related age signature for breast cancer.

We also observed clusters with *BRCA1/BRCA2* mutations and methylation (clusters 3 & 7, n=73 & 37), as previously described^13^. These tumours typically exhibited an HRD phenotype, and had elevated activity of the HRD-associated SNV signature (SNV-1). Cluster 3 was associated with SV-4 and SV-6 (tandem duplications), and more *BRCA1* and *PTEN* driver mutations than expected by chance. In contrast to clusters 1&2, patients in cluster 3 also tended to be younger than patients in other clusters. As expected, cluster 3 patients were predominantly from the Basal PAM50 class. Cluster 7 was associated with SV-5 (small deletions), loss of *BRCA2*, and Luminal B cancers. The majority (76%) of *BRCA1/2* cases fell into clusters 3&7, although *BRCA1/2* mutant tumours that fell outside these clusters often showed similar patterns of HRD-associated signature activities, albeit with increased activity of unrelated signatures (e.g. SV-3 in cluster 4). Of patients predicted by HRDetect^3^ to harbour HRD, 85% fell within the *BRCA1/2* (cluster 3&7) groups, demonstrating that the MMCTM output provides a substrate upon which known biological clusters are recovered, with further stratification as a result of SNV and SV integration.

Cluster 4 (n=67) had the highest activity of SV-3 (deletions and small inversions) and also contained more *TP53* mutant tumours than expected by chance alone. As this group contained examples of Her2, luminal A, luminal B and Basal PAM50 expression classes, we suggest cluster 4 represents an important group of 12% of breast cancers which transcend known molecular subtypes. Two clusters (clusters 5 & 6, n=53 & 41) were enriched for APOBEC signature activity (SNV-4 & SNV-6). Both of these APOBEC clusters were also enriched for HER2-positive tumours, and cluster 6 was enriched for *PIK3CA* driver mutations, relating Her2-amplification and APOBEC deamination processes for approximately 17% (cluster 5 + cluster 6) of breast cancers, as previously reported^25^. Additional small groups (Cluster 8 (n=18) and Cluster 9 (n=8)) contained tumours enriched for tandem duplications (SV-8) and association with defective DNA mismatch repair (MMRD, see COSMIC 6, 15, 20, 26), and SV-3 (deletions and small inversions), respectively, and consistent with previous reports^26^.

### SNV and SV signature activity segregates ovarian cancer cases into prognostically distinct groups

A recent analysis of ovarian tumours revealed a novel high-grade serous carcinoma (HGSC) sub-group with relatively worse prognosis, characterized by increased frequency of fold-back inversions (FBI)^2^. Their analysis combined NMF-based SNV signature analysis with ad-hoc SV and copy number variant (CNV) features. Here we expanded on some of their findings using the MMCTM on a merged data set consisting of 133 cases from Wang et al.^2^ and 62 cases from the ICGC ovarian cancer whole genome dataset^27^.

We fit 6 SNV and 7 SV signatures to mutation counts from the 195 ovarian cancer genomes (Fig. 3a,b, Supplementary Fig. 8, Supplementary Data 4, see Methods for signature count selection), including endometrioid carcinomas (ENOC), clear cell carcinomas (CCOC), granulosa cell tumours (GCT), and HGSC (Supplemental Table 1). Amongst the resultant SNV signatures were the previously described HRD (SNV-1, COSMIC 3), MMRD (SNV-2 & SNV-4, COSMIC 6, 15, 20, 26), APOBEC (SNV-3, COSMIC 2 & 13), *POLE* (SNV-5, COSMIC 10), and age-related signatures ((SNV-6, COSMIC 1), Supplementary Fig. 5, see also for a comparison to the breast SNV signatures). The SVs included signatures for tandem duplications (SV-1, SV-3, SV-6), translocations (SV-2), small deletions (SV-4), intra-chromosomal SVs generally >1Mbp (SV-5), and FBI and deletions (SV-7). The association of deletions with FBI can be understood in terms of the underlying cause of FBI: breakage-fusion-bridge cycles. After the loss of a telomere, sister chromatids fuse and are then pulled apart during mitosis, producing one chromosome with a foldback-inversion and another with a terminal deletion.

**Figure 3:**
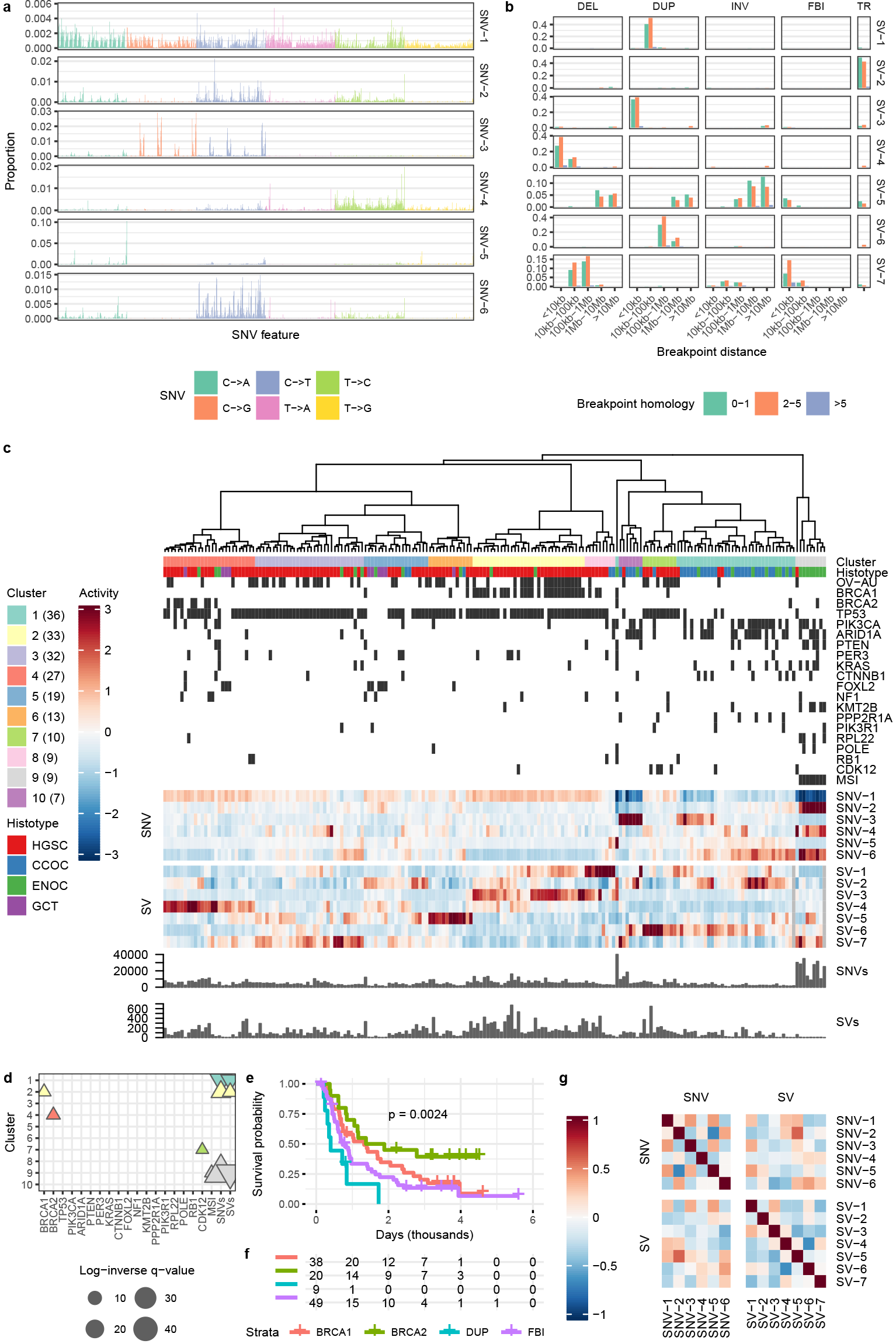
Ovarian cancer mutation signature analysis. **a** SNV mutation signatures. SNVs are organized according to the SNV type (color). Within each type, SNVs are further organized into the pattern of flanking nucleotides (AA-AA, AA-AC, … TT-TG, TT-TT). **b** SV mutation signatures. SVs are grouped by type (DEL: deletion, DUP: tandem duplication, INV: inversion, FBI: foldback inversion, TR: translocation). **c** Heatmap of relative signature activities in ovarian cancer samples. Each heatmap column represents a single case, and is composed of the proportions of SNV and SV signatures output from the MMCTM model. The values for each signature (row) have been standardized. Heatmap display has been truncated to ±3. Cases have been heirarchically clustered according to these transformed signature prevalences and cluster labels are indicated with colors underneath the dendrogram. The number of cases in each cluster is indicated in paretheses in the cluster legend. Cases from the ICGC OV-AU project are indicated with black bars, as is microsatellite instability (MSI) and gene mutation status. Samples with zero mutations for a mutation type also have greyed mutation signature activity cells. The number of SNVs for the POLE mutant case has been truncated to 40k in the barplot; The actual number is 596,135. d Annotation associations for sample clusters. Upward- and downward-pointing triangles indicate enrichment and depletion, respectively. Colors correspond to cluster colors indicated in the heatmap. **e** Kaplan-Meier curves for HGSC samples only. **f** Risk table for HGSC samples only. Kaplan-Meier curve plots and risk tables share x-axes. **g** Correlation heatmap between SNV and SV signatures.

We clustered the tumours according to their joint SNV and SV signature prevalences, which resulted in 10 groups (Fig. 3c, Supplementary Data 6, 7). Cluster 1 (n=36) contained mainly CCOC and ENOC tumours enriched for the age-associated signature (SNV-6), translocations (SV-2), and duplications (SV-6, see Supplementary Fig. 7, Supplementary Data 8 for cluster-signature associations). While the original study identified one HRD signature (SNV-1) group^2^, our analysis here produced two major HRD clusters (2 & 4, n=33 & 27), roughly defined by tumours with tandem duplications (SV-1 & SV-3) coupled with loss of *BRCA1* (see Fig. 3d, Supplementary Data 9 for cluster-annotation assocations), and small deletions (SV-4) coupled with loss of *BRCA2*, respectively. Cluster 2 also had greater activity of SVs compared to other ovarian tumours. Cluster 8 (n=9) included some *BRCA1* mutated tumours, and is distinct from cluster 2 due to greater enrichment of tandem duplication signature SV-1. The association of *BRCA1/2* status with tandem duplication and deletion SV signatures has been reported in breast cancer tumours^13^, and was reflected in our analysis of the 560 breast cancer dataset (Fig. 2, described above), providing strong evidence for BRCA1-like and BRCA2-like HRD sub-strata crossing tumour types.

Cluster 3 (n=32) was associated with enrichment of FBI (SV-7), cluster 5 (n=19) with translocations (SV-2), and cluster 6 (n=13) with large intra-chromosomal rearrangements (SV-5). Cluster 5 & 6 also included low-level FBI signature activity. Cluster 7 (n=10) was associated with higher activity of the duplication signature SV-6, and *CDK12* mutations, an association supported by a previous study^28^. Clusters 2-8 comprised mainly HGSC tumours, although each of these clusters also included tumours of other histotypes. For example, clusters 4-6 include GCT tumours. Cluster 9 (n=9) includes all microsatellite instable (MSI) ENOC tumours, and was associated with a mismatch repair deficient signature (SNV-2), the age-related signature (SNV-6), and higher numbers of SNVs. Cluster 10 (n=7) included 7 tumours highly enriched for APOBEC signature (SNV-3) activity.

By inspecting the signature correlations output by the MMCTM model (Fig. 3g, Supplementary Data 5), we saw that the HRD SNV signature (SNV-1) was positively correlated with the small tandem duplication signature SV-1 (r=0.29), as may be expected from the underlying biology of these signatures. The age-related signature (SNV-6) is positively correlated with SV signatures SV-5 (r=0.31) & SV-6 (r=0.41), which represent large intra-chromosomal SVs and duplications, respectively. The age-related and large intra-chromosomal SV signatures were also correlated in the breast cancer dataset analysis above. SNV-2 (MMRD-associated) is also positively correlated (r=0.59) with SV-5.

HGSC patient groups, defined by their mutation signature prevalences, differed in survival rates. We defined 4 HGSC super-clusters (see Methods), representing BRCA1-mutant (clusters 2 & 8, n=38), BRCA2-mutant (cluster 4, n=20), FBI (clusters 3, 5, 6, n=49), and tandem-duplicator tumours (cluster 7, n=9). We compared overall-survival amongst the HGSC super-clusters using the Kaplan-Meier method (Fig. 3e,f). The BRCA2/deletion cluster had the highest survival rate, while the tandem-duplicator group had the worst. Comparing the HGSC clusters in a pairwise fashion, the tandem-duplicator group had worse survival than the *BRCA1* group (adjusted log-rank p-value <0. 05) and the *BRCA2* group (adjusted log-rank p-value < 0.01). The FBI group had worse survival than the *BRCA2* group (adjusted log-rank p-value < 0.05). The BRCA1/tandem-duplication group had an intermediate survival rate, but the survival curve was not significantly different than those of the FBI or *BRCA2* groups (adjusted log-rank p > 0.05). While FBI was previously identified as a marker for poor prognosis^2^, activity of a mutational process linked with loss of *CDK12* and producing 100kbp-1Mbp tandem duplications appears to indicate even worse outcomes. Overall, the MMCTM analysis represented a refinement of signature-based prognostic stratification in HGSC indicating BRCA2-like HRD as the best performing group of patients, followed by BRCA1-like HRD, FBI and tandem duplicators.

## Discussion

Our results uncover a new landscape of mutational signatures in breast and ovarian cancers. Through principled, integrated inference and analysis of SNV and SV mutation signatures, our results reveal at once correlated signatures and novel disease sub-groups within DNA repair deficient tumours. Our findings have several implications for the field. The use of structural variations in signature analysis is less common than for point mutations, in part due to the relative paucity of whole-genome sequencing datasets. Here, we show the significant new value from their joint-interpretation, and set the framework for their simultaneous consideration across a broad range of tumour types.

This is evident through joint SNV & SV signatures-based subgroup identification in breast and ovary cancers, reproducing the association of tandem duplications within BRCA1-like and interstitial deletions within BRCA2-like cancers in two independent cancer types, with data from two independent studies. In the ovarian cancer cohort, this represents an important refinement in signatures-based tumour stratification, and furthermore we show how this has prognostic implication, superceding what could be derived from gene-based biomarkers (i.e. if only BRCA1 and BRCA2 mutation status were considered).

We have introduced a new formalism for mutation signature analysis in cancer genomes. Our approach models the correlation between signature activities, which improves method performance. Correlated topic models are significantly more robust to reduced mutation burden, which can occur in a number of scenarios. We have already described that signature extraction from SVs, at the level detected in the breast and ovarian datasets analysed here, benefits from correlated signature modeling. Analysis of other low-count mutation types may also benefit, for example mutations called from exome or single-cell sequencing experiments. Importantly, the statistical framework of the MMCTM is flexible and extensible. While here we show the advantage of integrated SNV and SV analysis, the MMCTM can seamlessly integrate other count-based features such as copy number events, double strand breaks, and telomeric insertions. As the field develops, we suggest a robust and extensible framework will be required to encode and integrate multiple feature types of the genome as they relate to mutational processes. The advantage of our relatively simple SNV and SV integration is evident and motivates further advances through multi-modal statistical modelling leading to richer biological interpretations of endogenous and potentially exogenous processes. In conclusion, our findings reinforce the importance of an integrated, holistic view of multiple classes of genomic scarring to drive discovery and characterization of mutation processes across human cancers.

## Methods

### Mutation data processing

Nucleotides flanking SNVs were extracted from human reference GRCh37. The number of each type of SNV (e.g. C → T) with a particular flanking sequence was counted. SV calls were split according to type (deletion, tandem duplication, inversion, foldback-inversion, translocation), the level of homology (0-1, 2-5, >5 bp), and breakpoint distance (<10kbp, 10-100kbp, 100 kbp-1Mbp, 1-10Mbp, >10Mbp), then counted. Foldback inversion calls were not included in the breast cancer dataset. Breakpoint distance bins are those used in a previous study on SV signatures^13^. Breakpoint distance was not calculated for translocations, as the concept is not applicable for this class of SVs. SNV and SV counts per sample were computed from the mutations counts used for signature analysis. Additional ovary sample gene mutation annotations were computed from SNV and indel calls according to the original paper.

Independent multi-modal correlated topic models The independent-feature multi-modal correlated topic model is based on a previously described independent mutation feature model^16^, as well as the multi-modal/field topic model extension to correlated topic models^21,22^(Fig. 1a). This model incorporates mutation types with contextual features unique to that type. The generative process is as follows:

Let

- *D* denote the number of documents
- *M* denote the number of modalities
- *K* denote the number of topics
- *N* denote the number of words in a document modality
- *I* denote the number of features in a modality’s words
- *J* denote the number of values in a feature

Then,

1. for each feature, *i*, in each topic, *k*, in each modality, *m*, draw 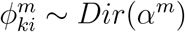
2. for each document, *d*, draw *η*^*d*^ ~ *N*(*μ*, Σ), where *η*^*d*^ is the concatenation of *η*^*dm*^ for all modalities, *i.e. η*^*d*^ = *η*^*d*1^,…, *η*^*dM*^
3. for each word, *n*, in each modality, *m*, draw word topic,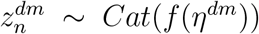, where 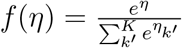
4. for each feature, *i*, in each word, *n*, above, draw 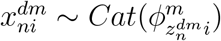

For simplicity, assume the number of topics, *K*, features, *I*, words, *N*, are the same across modalities and documents. Then the model likelihood is

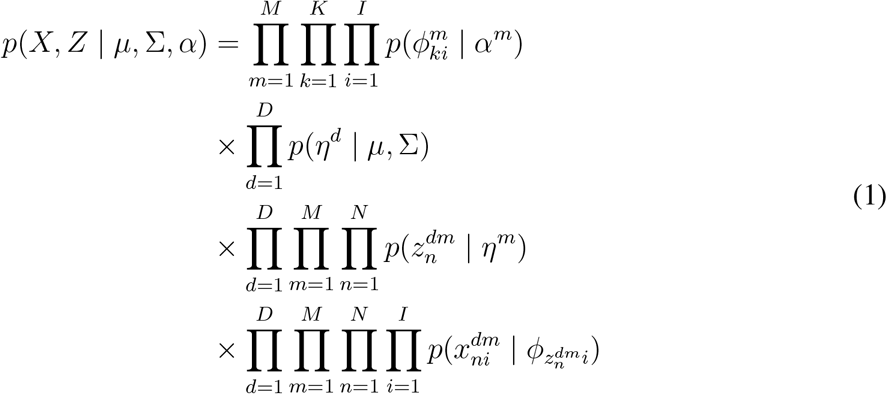

### Inference in topic models and NMF

For LDA and ILDA, parameters were inferred using mean-field variational bayes. For CTM, MMCTM, ICTM and IMMCTM, parameter inference was performed using mean-field variational EM. The MMCTM updates can be found in Salomatin et al.^22^. IMMCTM updates are similar, with modifications to allow for the independent feature construction of the mutation words.

The factorized mean-field variational Bayesian approximation for the IMMCTM is

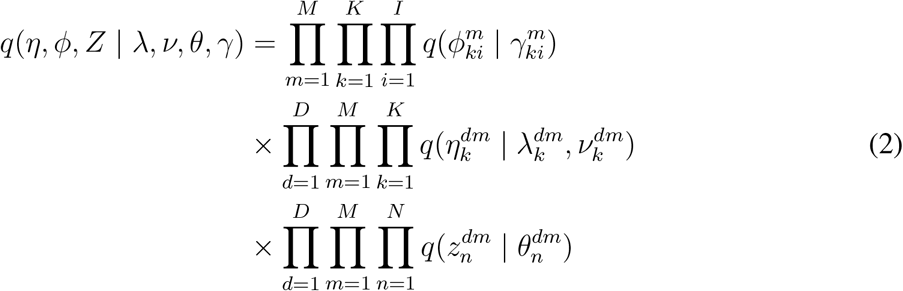

where

- 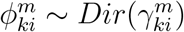
- 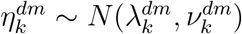
- 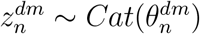
- The update for 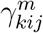 is

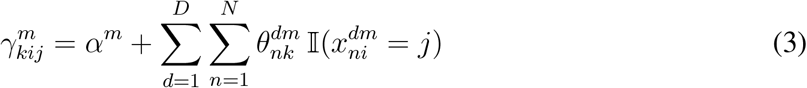

And the update for 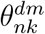 is

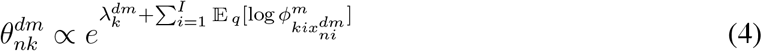

Single-modal correlated model parameters (CTM, ICTM) were inferred using MMCTM and IMMCTM, but with counts from a single mutation type. The probabilistic topic models were implemented similarly using the Julia language^29^. NMF models were fit using the coordinate descent solver implementation in the Scikit-learn library^30^ v0.18.1.

### Method comparison

NMF was run on both raw and normalized mutation counts. Normalization was performed by dividing mutation counts by sample totals, for each mutation type.

For log-likelihood-based comparisons, mutation counts were split according to a stratified 10 × 5 cross validation scheme; For each histotype, cases were split into 5 training and test sets. The splitting procedure was performed 10 times, resulting in 50 training and test sets.

Each method was run on each training set and evaluated on each corresponding test set, using random initialization. Evaluation was performed by randomly splitting the mutations in each test case into observed and hidden sets. Signature proportions for each test case were estimated using the observed test mutation counts, then the per-word log predictive likelihood was computed using the hidden test mutation counts. Methods were tested over a range of 2-12 signatures, as well as over a range of count subsets. Multi-modal topic models were given the same number of signatures for SNVs and SVs.

Count subset comparisons were performed by removing mutations from each genome, retaining only a given fraction. Mutations were randomly selected according to their type (e.g. AC(C→T)TT) and relative type proportions. These mutations were removed and the genome mutation counts updated. The updated mutation counts were then input to the compared methods. SNVs were subset to 1, 5, 10%, while keeping SVs at 100%. SVs were subset to 10, 15, 20%, while keeping SNVs at 100%. For the breast cancer dataset, the number of SNV and SV signatures was fixed at 5, the optimal number of SV signatures for NMF, and an SNV signature plateau-point for NMF run on raw counts.

Log predictive likelihoods were computed on test sets with signatures for SNVs and SVs separately. The likelihood computation involves the signature-mutation proportions fit with the training data, case-signature proportions estimated using the observed test counts, and the hidden test counts. The average per-word predictive log likelihood for a particular mutation type is given in equation 5.

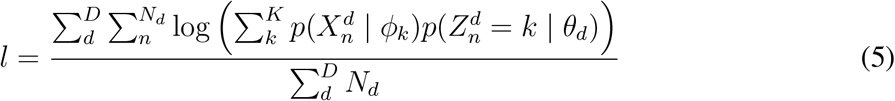

where *D* is the number of cases, *N*_*d*_ is the number of mutations in case *d*, *K* is the number of signatures, *X* is the mutations in case *d*, *Z* is the mutation-signature indicators, *ϕ*_*k*_ is the signature-mutation distribution, and *¸*_*d*_ is the document-signature distribution.

For the logistic regression classifier-based comparisons, each signature detection method was trained 10 times with 2-10 signatures, using the full 560 breast cancer dataset. For multi-modal methods, the same number of SNV and SV signatures was given. The sample-signature distributions were used as training data for the classifier along with previously published HRDetect-derived labels. Three types of tests were performed: using only SNV, only SV, or both SNV and SV sample-signature distributions. Stratified 5-fold cross-validation was performed for each test, resulting in 5 × 10 = 50 scores for each method, training data type, and setting of the number of signatures. The output score of cross validation is the mean accuracy of the logistic regression classifier. Parameter inference was performed using the Scikit-learn^30^ v0.18.1 implementation with the liblinear solver and maximum 10,000 iterations.

### Choosing the number of signatures

The number of signatures to estimate was selected using the cross validation scheme described in the method comparison. Log-likelihood values were plotted across 2-20 signatures and the elbow method was used to select the number of signatures (Supplementary Fig. 10).

### Fitting MMCTM to cancer datasets for downstream analysis

The model was initially fit to each dataset 1000 times for a limited number of iterations. *α* hyper-parameters were set to 0.1. Each restart is run until the relative difference in log predictive likelihood on the training data was < 10^−4^ between iterations. The restart with the best mean rank of the SNV and SV log predictive likelihoods was selected for fitting to convergence with a tolerance of 10^−5^.

### Case hierarchical clustering

Cases were clustered using case-signature proportions for SNV and SV signatures together. Proportion values were converted to Z-scores for each signature across cases. By standardizing the proportion values, the inter-case differences of low-prevalence signatures are given increased emphasis relative to higher-prevalence signatures. Hierarchical agglomerative clustering was performed using the euclidean metric, and Ward linkage. Discrete clusters were formed using the R dynamicTreeCut package^31^ v1.63 with `method=“hybrid”, deepSplit=FALSE`, and `minClusterSize=3`.

### Sample cluster enrichment and depletion tests

Enrichment of a sample cluster’s signature activity was tested using an unequal variance one-sided t-test against the signature activities of other clusters.

For the breast cancer dataset, cluster associations with ER, PR, HER2, MMRD, and PAM50 status were performed with a two-tailed Fisher’s exact test. Differences in Age or the number of SNVs and SVs were tested with two-tailed unequal variance t-tests. Driver gene mutation and HRDetect prediction associations were computed using a blocked permutation test.

The permutation tests were performed as follows: For each cluster, “new” clusters were generated by sampling cases without replacement from the full dataset. New clusters maintained the same ER, PR, and HER2 status composition as the original cluster. The difference in proportions of cases with the annotation of interest between the new cluster and all other cases was computed. Two-tailed p-values were calculated using equation 6:

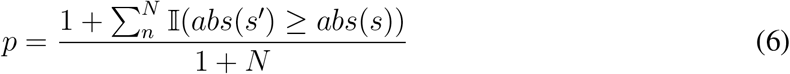

where *N* is the number of permutations (generated clusters), and *s* is the statistic of interest for the original cluster (*e.g.* difference in proportions of samples with loss of TP53), and *s′* is the same statistic for a generated cluster. This procedure attempts to correct for correlations between the tested annotations and ER, PR, and HER2 status.

Gene mutation status and MSI cluster associations in ovarian cancer were tested with the blocked permutation test described above, accounting for histotype rather than ER, PR, and HER2 status. Differences in SNV and SV counts were performed with two-tailed unequal variance t-tests. Due to the presence of a POLE mutant sample with a very high number of SNVs, t-tests for this statistic were performed on count ranks. The unequal variance t-test on ranked data is a robust alternative to Student’s t-test and the Mann-Whitney U test when assumptions are violated^32^.

Cluster-signature and cluster-annotation p-values within each dataset were corrected using the Benjamini & Hochberg method^33^.

### Survival analysis

HGSC cases grouped according to the hierarchical clustering were compared by estimating overall-survival Kaplan-Meier curves for each cluster, using the R survival package. Clusters 3, 5, and 6 were grouped as they are related in the hierarchical clustering, and had no significant difference in survival outcome. We call this the “FBI” group, due to higher activity of the FBI signature among HGSC cases. Similarly, clusters 2 and 8 form the “BRCA1” group. P-values were calculated using the log-rank test. Pairwise survival curve comparison p-values were adjusted using the Benjamini & Hochberg method^33^ implemented in the R `p.adjust` function.

## Code availability

Topic model code is available in a GitHub repository:
https://github.com/funnell/MultiModalMuSig.jl

## Data availability

Mutations and sample annotations for the 560 breast cancer landscape study^13^ were downloaded from the ICGC DCC (project BRCA-EU, https://dcc.icgc.org/releases). Additional sample annotations were obtained from related study supplementary files^3,13, 26^. Ovary mutation calls and sample annotations were obtained from Wang et al.^2^.

## Acknowledgements

We wish to acknowledge the generous long term funding support from BC Cancer. SPS is a Michael Smith Foundation for Health Research (MSFHR), holds a Canadian Institutes for Health Research (CIHR) Foundation grant and holds Canada Research Chairs. The authors wish to acknowledge the funding support from the Discovery Frontiers: Advancing Big Data Science in Genomics Research program (grant no. RGPGR/448167-2013, ‘The Cancer Genome Collaboratory’), which is jointly funded by the Natural Sciences and Engineering Research Council of Canada, the Canadian Institutes of Health Research, Genome Canada, and the Canada Foundation for Innovation, and with in-kind support from the Ontario Research Fund of the Ministry of Research, Innovation and Science.

## Author contributions

T.F. developed the algorithms and software, performed all data analysis and wrote the manuscript. A.Z. edited the manuscript.

Y.J.S. and R.L. provided assistance with cancer data processing.

D.G. provided data analysis and software development support.

S.M. assisted with statistical analysis.

A.B. supported software development and data analysis.

Y.K.W. supported the ovarian cancer data analysis.

P.C.B. supported data interpretation and edited the manuscript.

S.P.S. provided project conception and oversight and is the senior responsible author.

